# Hi-Cformer enables multi-scale chromatin contact map modeling for single-cell Hi-C data analysis

**DOI:** 10.1101/2025.08.04.668453

**Authors:** Xiaoqing Wu, Xiaoyang Chen, Rui Jiang

## Abstract

Single-cell Hi-C captures the three-dimensional organization of chromatin in individual cells and provides insights into fundamental genomic processes such as gene regulation and transcription. While analyses of bulk Hi-C data have revealed multi-scale chromatin structures like A/B compartments and topologically associating domains, single-cell Hi-C data remain challenging to analyze due to sparsity and uneven distribution of chromatin contacts across genomic distances. These characteristics lead to strong signals near the diagonal and complex multi-scale local patterns in single-cell contact maps. Here, we propose Hi-Cformer, a transformer-based method that simultaneously models multi-scale blocks of chromatin contact maps and incorporates a specially designed attention mechanism to capture the dependencies between chromatin interactions across genomic regions and scales, enabling the integration of both global and fine-grained chromatin interaction features. Building on this architecture, Hi-Cformer robustly derives low-dimensional representations of cells from single-cell Hi-C data, achieving clearer separation of cell types compared to existing methods. Hi-Cformer can also accurately impute chromatin interaction signals associated with cellular heterogeneity, including 3D genome features such as topologically associating domain-like boundaries and A/B compartments. Furthermore, by leveraging its learned embeddings, Hi-Cformer can be extended to cell type annotation, achieving high accuracy and robustness across both intra- and inter-dataset scenarios.

## Introduction

The 3D organization of chromatin plays a vital role in regulating genome function^1^. By shaping the spatial proximity between regulatory elements and genomic loci, 3D genome structure facilitates the transcriptional regulation^2^. Moreover, the 3D organization of chromatin is closely associated with DNA replication timing^3,4^, and contributes to the functional compartmentalization of the nucleus^5^. Disruptions in 3D genome structure have been shown to rewire regulatory architectures and cause pathogenic gene expression ^6–8^, underscoring the functional relevance of chromatin organization in the cellular context.

Chromosome conformation capture (3C) techniques^9^, such as Hi-C^10^ have significantly advanced the study of chromatin conformation, enabling the identification of 3D genome structures such as A/B compartments^5^, topologically associating domains (TADs)^11–13^, and chromatin loops^14^. In recent years, single-cell Hi-C (scHi-C) has enabled the investigation of 3D chromatin organization at the single-cell level, providing valuable insights into chromatin interactions that are associated with cell heterogeneity^15–19^. However, unlike other single-cell sequencing data that are typically represented as a feature vector per cell, scHi-C produces a series of matrices of sparse contact maps with pronounced genomic-distance-dependent bias, leading to strong diagonal signals and complex multi-scale local patterns that pose challenges for downstream analysis^20^.

Computational methods have been developed to address the challenges posed by scHi-C data and facilitate its analysis^21,22^. Non-deep learning methods typically adopt graph-based or statistical models as analytical frameworks^23–26^. For instance, random walk with restart (RWR) and its variants are widely used for imputing contact maps, with scHiCluster^23^ being a representative example. Statistical methods such as principal component analysis (PCA) and latent Dirichlet allocation (LDA) are applied to obtain low-dimensional representations of cells^23,24^. Recent advances in deep learning have enabled the development of methods that capture the associations between complex cellular heterogeneity and chromatin interactions^27–30^. For example, Higashi^27^, a representative self-supervised method, used hypergraph representation learning for the embedding and imputation of scHi-C data. scVI-3D^28^ employed variational autoencoders to perform embedding and denoising of scHi-C data. scDEC-Hi-C^29^ introduced a generative adversarial network (GAN)-based framework. scHiClassifier^30^, a supervised method, was also developed for annotating cell types with scHi-C data.

Nevertheless, most existing methods model either entire chromosomal contact maps or pairwise interactions at a fixed resolution, without explicitly capturing the complex multi-scale structures present in chromatin contact matrices—structured patterns of varying sizes that carry cellular heterogeneity information about 3D genome organization^21^. This limitation hinders the simultaneous analysis of global and multi-scale local chromatin interaction features, restricting the ability of these methods to capture comprehensive chromatin interaction patterns.

To address these challenges, we propose Hi-Cformer, a transformer-based method that represents chromatin contact maps of a cell as sequences of embeddings, analogous to the way language models encode sequences of words. The model integrates local structural features derived from multi-scale contact map blocks with global contextual information from entire chromosomal maps. Hi-Cformer employs a chromosome-aware hierarchical attention mechanism that captures intra-chromosomal dependencies of multi-scale interaction and enables effective integration of inter-chromosomal context. This architecture preserves both fine-grained and overall chromatin interaction patterns, enabling comprehensive modeling of 3D genome organization similar to how language models capture contextual relationships within sequences.

The above modeling design of Hi-Cformer supports its strong performance on multiple downstream tasks. Hi-Cformer can generate informative cell representations from sparse and complex scHi-C data, capturing cellular heterogeneity and enabling accurate cell-type discrimination across datasets with varying sequencing protocols, depths, and cell type compositions. Beyond dimensionality reduction, Hi-Cformer achieves more reliable imputation of sparse scHi-C contact maps than existing methods, by capturing cell-type-specific chromatin patterns while retaining single-cell-level heterogeneity. Hi-Cformer enhances the detection of cell-type-specific chromatin features, such as TAD-like domain boundaries and A/B compartments, and facilitates the analysis of cell-type heterogeneity in chromatin organization. As a flexible and extensible framework, Hi-Cformer can be adapted for supervised cell type annotation, showing strong performance in both intra- and inter-dataset scenarios. Together, these capabilities position Hi-Cformer as an effective and versatile tool for single-cell 3D genome analysis.

## Results

### Overview of Hi-Cformer

Hi-Cformer takes the intra-chromosomal contact maps of all chromosomes from a single cell at a given resolution as input, and outputs reconstructed contact maps at three hierarchical scales: a cell-level map formed by concatenating all intra-chromosomal maps, chromosome-level maps for individual chromosomes, and block-level submaps with varying sizes. To achieve this objective, Hi-Cformer represents all intra-chromosomal contact maps of a cell as an ordered sequence of embeddings, following the input representation paradigm used in language models, thereby enabling the incorporation of both block-level and global-context 3D chromatin features while retaining cell-level resolution. Hi-Cformer consists of three modules: the multi-scale encoder module, the transformer module, and the multi-scale decoder module (Fig. 1).

**Fig. 1.**
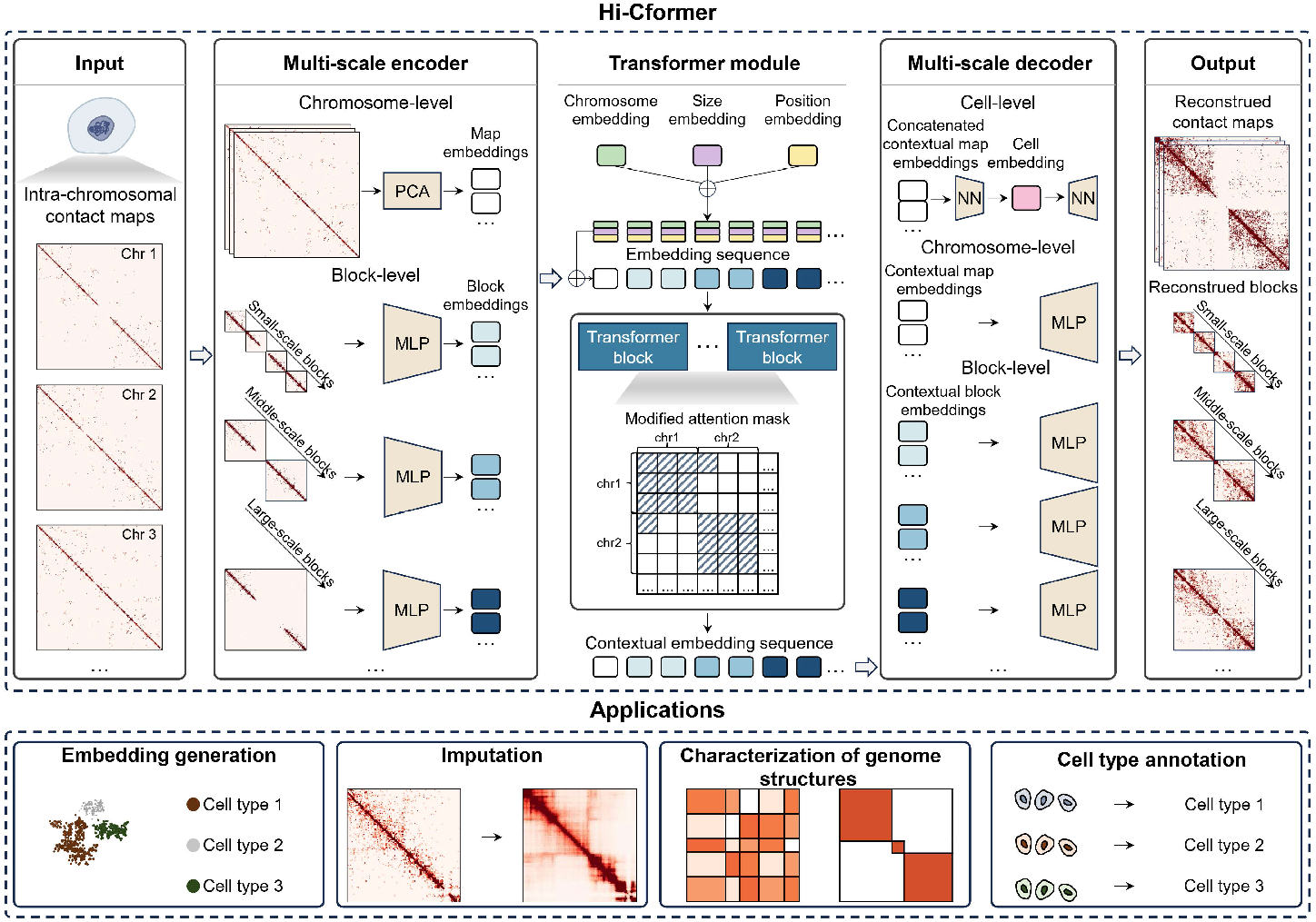
Overview of Hi-Cformer architecture and applications. For self-supervised model training, the single-cell intra-chromosomal contact maps are fed to Hi-Cformer. The model first embeds both intra-chromosomal contact maps and blocks at multiple scales using PCA and dedicated MLPs, generating a comprehensive embedding sequence for each cell. The embedding sequence is added to chromosome, size, and position embeddings, and then passed through transformer blocks with a modified attention mask that enables the model to capture both intra- and inter-chromosomal dependencies. The contextual embeddings are decoded at three levels to reconstruct genome contact patterns: a cell-level reconstruction that outputs a unified whole-cell representation, chromosome-level reconstructions that capture whole-cell contact information via per-chromosome decoding, and block-level reconstructions that recover local structures at multiple scales. Hi-Cformer facilitates embedding generation, imputation, characterization of 3D genome structures and cell type annotation of scHi-C data.

The multi-scale encoder module encodes diagonal blocks of varying sizes from the intra-chromosomal contact map of each chromosome into fixed-dimensional vectors, referred to as block embeddings, as these regions correspond to genomically proximal loci with denser and more informative interaction patterns, often reflecting domain-like chromatin structures such as TADs. These block embeddings, together with chromosomal map embeddings of all chromosomes obtained via PCA, are then organized into a sequence ordered by chromosome index, block size, and position. The resulting sequence plays a role analogous to token sequences in language models, enabling sequential modeling of chromatin contacts using a transformer.

The transformer module processes the output sequence from the multi-scale encoder, with additional embeddings for chromosome identity, block size, and genomic position added element-wise to each embedding in the sequence. These additional embeddings help the model distinguish among chromatin regions from different chromosomes, genomic positions, and scales. The resulting sequence is processed through transformer blocks^31^ with modified attention masks that constrain block-level attention within chromosomes, and enable global context integration through chromosomal map embeddings.

The multi-scale decoder module uses contextual block embeddings and chromosomal map embeddings derived from the output of the transformer module as input. These two types of embeddings are then fed into different multilayer perceptrons (MLPs) for three reconstruction tasks. The decoders produce cell embeddings as intermediate representations, along with multi-scale imputed contact maps as the outputs of these tasks.

Hi-Cformer is trained in a self-supervised manner to reconstruct input signals using a tailored masked language modeling (MLM) task^32^, which encourages the model to capture long-range dependencies. The flexible architecture of Hi-Cformer allows introducing new embeddings to impute contact maps for chromatin regions of different resolutions, while also supporting supervised tasks like cell type annotation through the incorporation of a discriminator.

### Hi-Cformer generates embeddings capturing rich cellular heterogeneity

Given the complex nature of raw scHi-C data, direct analysis of contact maps remains challenging. Therefore, obtaining low-dimensional representations of cells (referred to as cell embeddings) is essential for effective downstream analysis. To derive these embeddings, the contextual chromosomal map embeddings are concatenated into a whole-cell representation, which is then passed through a dimension-reduction and expansion process during cell-level reconstruction. The latent vector used for this reconstruction serves as the cell embedding, capturing the integrated multi-chromosomal structural information of each cell. We assessed the cell embeddings generated by Hi-Cformer on five scHi-C datasets^15–18^ with known cell type labels at 1-Mb resolution, and compared Hi-Cformer against six baseline methods: Higashi^27^, scDEC-Hi-C^29^, HiCRep/MDS^26,33^, scHiCluster^23^, PCA^34^, and LDA^24,35^. To quantitatively evaluate the ability of each method for revealing cellular heterogeneity, we used the Leiden algorithm^36^ to cluster the cell embeddings, with the number of clusters set to match the ground-truth labels by tuning the resolution parameter via binary search. We then used normalized mutual information (NMI) and adjusted Rand Index (ARI) scores as metrics to evaluate clustering performance, where higher values indicate a more accurate representation of cellular heterogeneity. Besides clustering-based metrics (NMI and ARI), we additionally incorporated a clustering-independent measure, the cell-type local inverse Simpson’s index (cLISI)^37^, which assesses the separation of cell types directly in the embedding space. Further details on the datasets, experimental setup, evaluation criteria, and baseline methods are provided in Methods.

As shown in the left panels of Fig. 2a–2e, the cell embeddings derived from Hi-Cformer outperform those from the baselines in the clustering task, with average improvements of 6.41% in NMI and 29.35% in ARI over the second-best method, Higashi. Together with the consistently strong cLISI scores across datasets (Supplementary Fig. S1–S5), these metrics collectively indicate that Hi-Cformer generates well-structured embeddings in which distinct cell types are clearly resolved. Importantly, these gains were observed across datasets with markedly different sequencing depths, noise levels, and cell-type compositions, highlighting the robustness of our method. We observed that non-deep learning methods, such as LDA, obtain substantial performance fluctuations across datasets. In contrast, Hi-Cformer demonstrates more stable and consistent performance. Closer examination of individual datasets further illustrates Hi-Cformer’s strengths. On the Ramani2017 dataset (Fig. 2a), Hi-Cformer not only achieves the highest NMI and ARI but also resolves the rare GM12878 population (33 cells), demonstrating its sensitivity to low-abundance cell types. On the Lee2019 dataset (Fig. 2b), Hi-Cformer distinguishes neuronal subtypes that were not separable using scHi-C data in the original study^16^, suggesting that it captures subtle chromatin features associated with neuronal lineage diversification. The results of the Tan2021A and Tan2021B datasets (Fig. 2c and Fig. 2d) indicate that Hi-Cformer effectively learns cell-type-specific chromatin signatures that generalize across complex brain datasets. On the higher-depth Wu2024 dataset (Fig. 2e), whose median raw contacts per cell is approximately 2.5 times that of the next deepest dataset, Hi-Cformer maintains its advantage, showing that it can effectively exploit the richer contact information provided by high-quality sequencing.

**Fig. 2.**
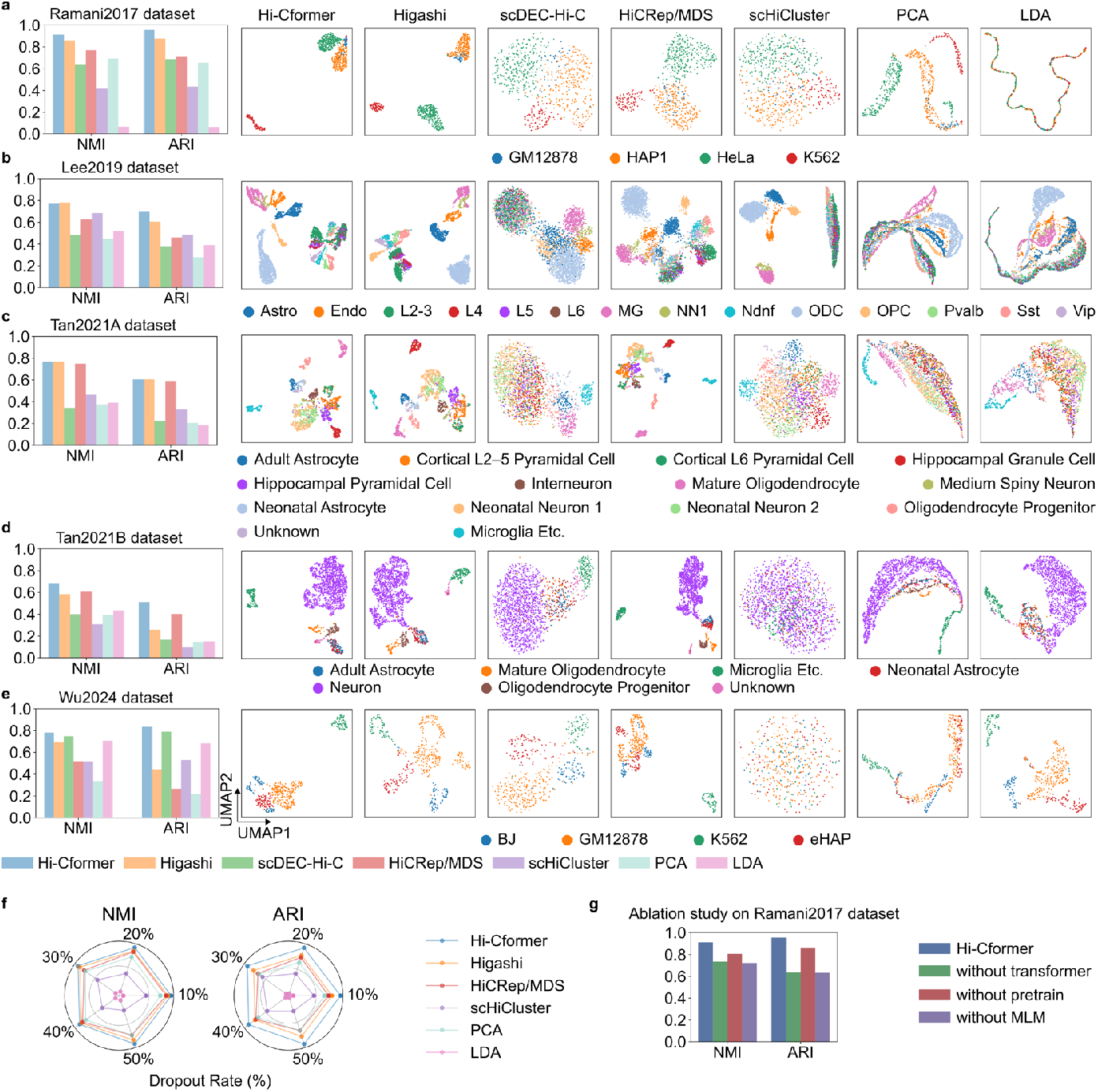
Evaluation and visualization of embeddings generated by different methods on scHi-C data. **a**–**e**, Quantitative evaluation of Hi-Cformer on the Ramani2017 (**a**), Lee2019 (**b**), Tan2021A (**c**), Tan2021B (**d**), and Wu2024 (**e**) datasets by comparing with six baseline methods: Higashi, scDEC-Hi-C, HiCRep/MDS, scHiCluster, PCA, and LDA. Cell embeddings were generated from contact maps at 1-Mb resolution and clustered using the Leiden algorithm with the number of clusters set to match the ground-truth labels. Left panels report clustering performance measured by normalized mutual information (NMI) and adjusted Rand index (ARI); right panels report UMAP visualization of the embeddings, colored according to ground-truth cell types. **f**, Robustness analysis on the Ramani2017 dataset under simulated dropout noise. **g**, Ablation analysis of model components on the Ramani2017 dataset. Comparison of performance under different ablation settings, i.e., omitting the transformer module, the preheating strategy, or the MLM task.

During the sequencing procedure, dropout events often occur, leading to high noise and sparsity in the scHi-C data. To assess the robustness of Hi-Cformer under such conditions, we tested its performance on the simulated dataset by randomly removing raw contacts from the Ramani2017 dataset. Across all five datasets with various dropout rates, Hi-Cformer consistently outperforms baseline methods, demonstrating its strong robustness to dropout events (Fig. 2f). To investigate the contributions of each component and training strategy in Hi-Cformer, we performed ablation experiments by omitting the transformer module, preheating strategy, and MLM task in the training stage, respectively. The results (Fig. 2g) show that removing the transformer module leads to a greater decline in embedding performance than removing preheating, as the transformer is essential for capturing intra- and inter-chromosomal dependencies of contact maps, while preheating mainly accelerates the training process. Notably, the models without the MLM task and those without the transformer module perform similarly, underscoring the critical role of the MLM task in effectively training the transformer module. Additionally, we explored the impact of selecting blocks of different sizes on embedding results. The results indicate that leveraging multi-scale blocks as input enhances embedding quality relative to single-scale designs (Supplementary Fig. S6). Together, these findings emphasize the critical role of each component in enabling Hi-Cformer to effectively model the 3D chromatin organization.

### Hi-Cformer provides reliable imputation of scHi-C contact maps

ScHi-C data are inherently sparse and noisy due to limited sequencing depth, motivating the need for computational imputation to recover biologically meaningful chromatin interaction patterns^21^. For imputation, Hi-Cformer uses its cell-level reconstruction output—a whole-cell contact map at the same resolution as the input. To evaluate whether Hi-Cformer effectively addresses these challenges, we first benchmarked its performance on the Ramani2017 dataset at 1-Mb resolution, using bulk Hi-C contact maps from Rao2014 dataset^14^ as an external reference. Compared with two established imputation methods, Higashi and scHiCluster, Hi-Cformer achieves a median improvement of 68.95% in Pearson correlation coefficient and 66.15% in cosine similarity over the second-best baseline (Higashi) across two cell types (Fig. 3a). When pseudo-bulk maps averaged from single-cell data of the same cell type were used as the reference, this comparison reflected the model’s ability to recover cell-type-specific chromatin structures. Under this evaluation setting, Hi-Cformer achieved median improvements of 70.60% in Pearson correlation coefficient and 68.15% in cosine similarity over Higashi (Fig. 3b). The advantages of Hi-Cformer remained consistent across the other datasets that we tested, as shown in Supplementary Fig. S7–S10, which indicated strong robustness across different experimental conditions and cell types. Importantly, these improvements are not merely numerical gains. scHi-C contact maps contain hierarchical structures spanning local domains to global organizations, and Hi-Cformer’s ability to accurately recover these features reflects its architectural design. Its multi-scale encoder explicitly extracts block-level embeddings corresponding to TAD-like structures, while chromosome-level embeddings preserve global context. These representations are jointly processed by a transformer with scale- and position-aware attention mechanisms, enabling the model to integrate local high-density signals with long-range chromosomal interactions.

**Fig. 3.**
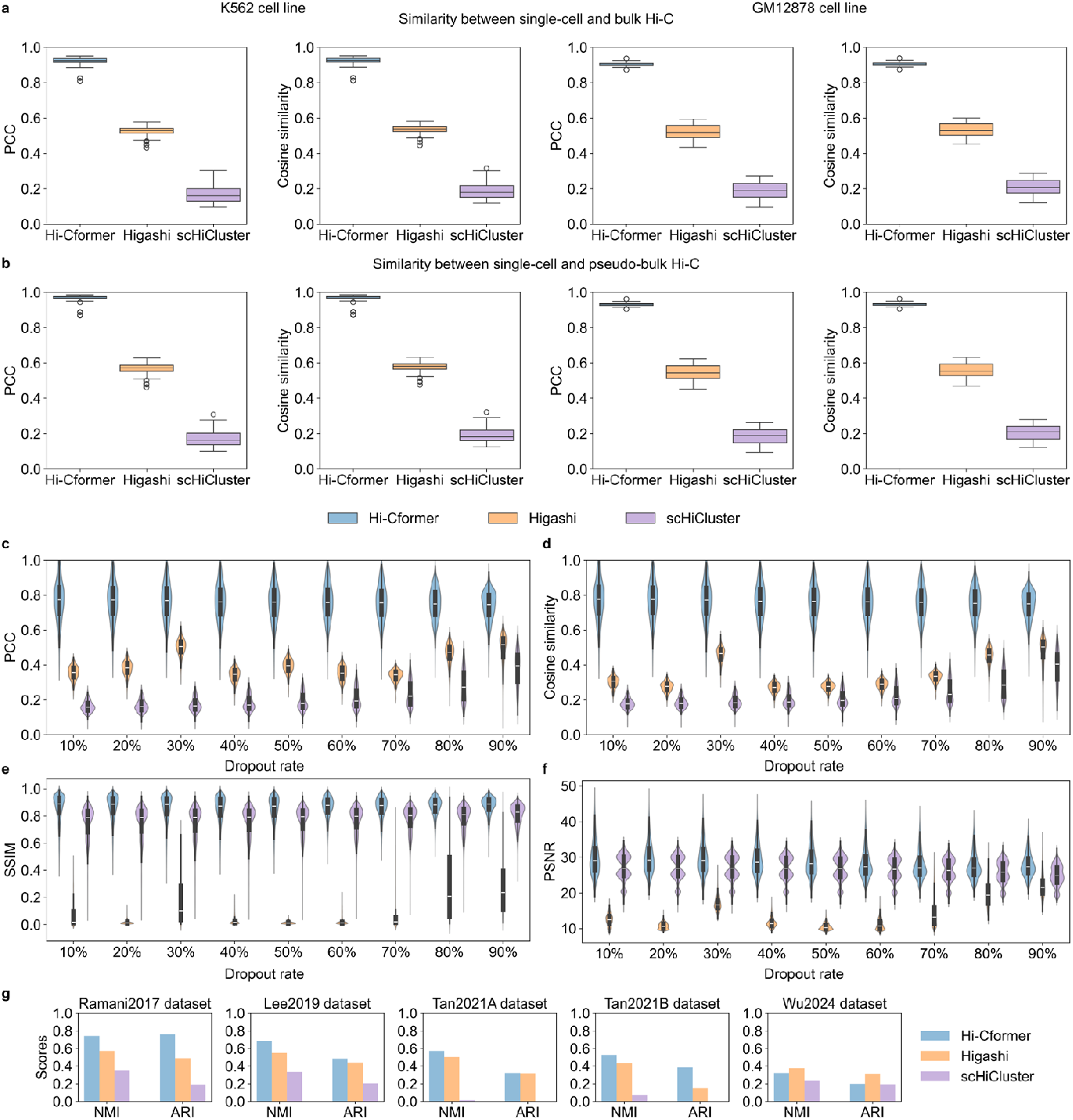
Comparison of Hi-Cformer with two baseline imputation methods from different evaluation perspectives. **a**, Pearson correlation coefficients (PCCs) and cosine similarities between scHi-C data imputed by three methods from the Ramani2017 dataset and corresponding bulk Hi-C data from Rao2014 dataset, for the K562 and GM12878 cell types (the left two subpanels in a correspond to K562, and the right two to GM12878). **b**, PCCs and cosine similarities between scHi-C data imputed by three methods from the Ramani2017 dataset and corresponding pseudo-bulk contact maps (aggregated from raw single-cell maps), for the K562 and GM12878 cell types (the left two subpanels in b correspond to K562, and the right two to GM12878). **c**, PCCs between imputed and ground truth data on simulated dropout datasets generated from the Ramani2017 dataset with varying dropout rates (10% to 90%). **d**, Cosine similarities between imputed and ground truth data on simulated dropout datasets generated from the Ramani2017 dataset with varying dropout rates (10% to 90%). **e**, SSIMs between imputed and ground truth data on simulated dropout datasets generated from the Ramani2017 dataset with varying dropout rates (10% to 90%). **f**, PSNRs between imputed and ground truth data on simulated dropout datasets generated from the Ramani2017 dataset with varying dropout rates (10% to 90%). In **a**–**f**, the lower, middle, and upper edges of the boxes denote the lower quartile, median, and upper quartile, respectively, and error bars denote the maximum and minimum values. **g**, NMI and ARI scores for clustering results after dimensionality reduction of imputed data by three methods on five benchmark scHi-C datasets.

Furthermore, to enable a quantitative evaluation of imputation accuracy at the level of individual cells, we benchmarked Hi-Cformer and baselines on five simulated datasets by introducing varying degrees of dropout events to the raw scHi-C contact maps. The raw contact maps before dropouts serve as ground truth, allowing for direct comparison of the ability of each method to reconstruct missing chromatin interactions. Hi-Cformer consistently outperforms Higashi and scHiCluster across all four evaluation metrics, including PCC, cosine similarity, structural similarity index measure (SSIM), and peak signal-to-noise ratio (PSNR), and also maintains stable performance across varying dropout rates, indicating high accuracy and robustness for imputation (Fig. 3c–3f). Higher SSIM values indicate that the imputed contact maps not only match the ground-truth interaction intensities but also preserve their spatial structural organization, including local patterns such as TAD-like domains, boundary sharpness, and regional contrast. This is particularly important for scHi-C, where biological information is often encoded in the relative arrangement of contact patterns rather than in absolute interaction counts alone. Likewise, higher PSNR reflects a lower level of reconstruction noise relative to true chromatin signals, demonstrating that Hi-Cformer effectively recovers genuine interaction patterns without introducing artificial smoothing or noise amplification. The lower concordance with ground truth obtained by Higashi and scHiCluster may be partially related to their tendency to generate denser contact maps during imputation, which may inflate apparent interaction frequencies but reduce concordance with true structure.

Finally, to assess whether imputation with Hi-Cformer can improve accuracy of downstream analyses, we applied PCA on contact maps imputed by different methods to reduce dimensionality to 64 principal components, followed by benchmarking the performance of cell clustering using corresponding metrics. The results demonstrated that imputation with Hi-Cformer can produce substantially more accurate and stable cell-type separation than baseline methods (Fig. 3g). This suggests that Hi-Cformer not only enhances pairwise contact recovery but also effectively preserves cell-type-defining chromatin organization patterns, enabling more reliable interpretation of cellular heterogeneity in scHi-C datasets.

### Hi-Cformer facilitates the identification of cell-type-specific structures

Cell-type-specific structures of chromatin contact maps have been a central focus of scHi-C analysis^38^. In this section, we explored how imputed scHi-C data with Hi-Cformer facilitates the identification of cell-type-specific structures such as TAD-like domain boundaries and A/B compartments. Prior scHi-C studies have shown that these structures can be detected from single-cell data, but their recovery is often hindered by sparsity and noise^23,27^. Leveraging its imputation output, Hi-Cformer produces contact maps from which compartment patterns and TAD-like boundaries can be more reliably inferred.

We first focused on a genomic region (chr9:132,950,000–138,000,000) previously reported to differ between the K562 and GM12878 cell types^29^. Within this region, the ABL1 gene locus (chr9:133,710,641–133,763,062) is of particular interest. ABL1 is a known oncogene involved in chronic myeloid leukemia (CML). While both BCR and ABL1 are present in K562 and GM12878 cells, only K562 harbors the BCR-ABL fusion gene, resulting from a translocation between chromosome 9 and 22^39^. This may contribute to the differences observed in the scHi-C contact maps of this region. To better resolve this region, we leveraged Hi-Cformer’s ability to encode contact map blocks of varying sizes into fixed-dimensional embeddings, which enables flexible resolution handling. Specifically, we introduced an additional embedding for the selected 50-kb resolution region, accompanied by a dedicated encoder and decoder, enabling Hi-Cformer to impute this high-resolution region (details in Methods). We conducted imputation experiments for this region on K562 and GM12878 cells from the Ramani2017 dataset and presented the resulting chromatin contact maps. For both cell types, raw and imputed scHi-C contact maps were averaged to generate pseudo-bulk data, which were then compared to the corresponding bulk Hi-C maps from Rao2014 dataset^14^. As shown in Fig. 4a, the imputed pseudo-bulk maps more clearly revealed structural differences (chr9:133,550,000–134,050,000) between K562 and GM12878 cells, particularly around the ABL1 locus. Within this genomic region, H3K27ac ChIP-seq signals are enriched in K562 cells compared to GM12878 cells (Fig. 4b), consistent with enhanced regulatory activity associated with the oncogenic fusion event. In addition to directly inspecting local structural differences, we further assessed large-scale chromatin organization by analyzing the merged scHi-C correlation matrices. Imputation with Hi-Cformer strengthened the checkerboard-like patterns indicative of A/B compartmentalization (Fig. 4c).

**Fig. 4.**
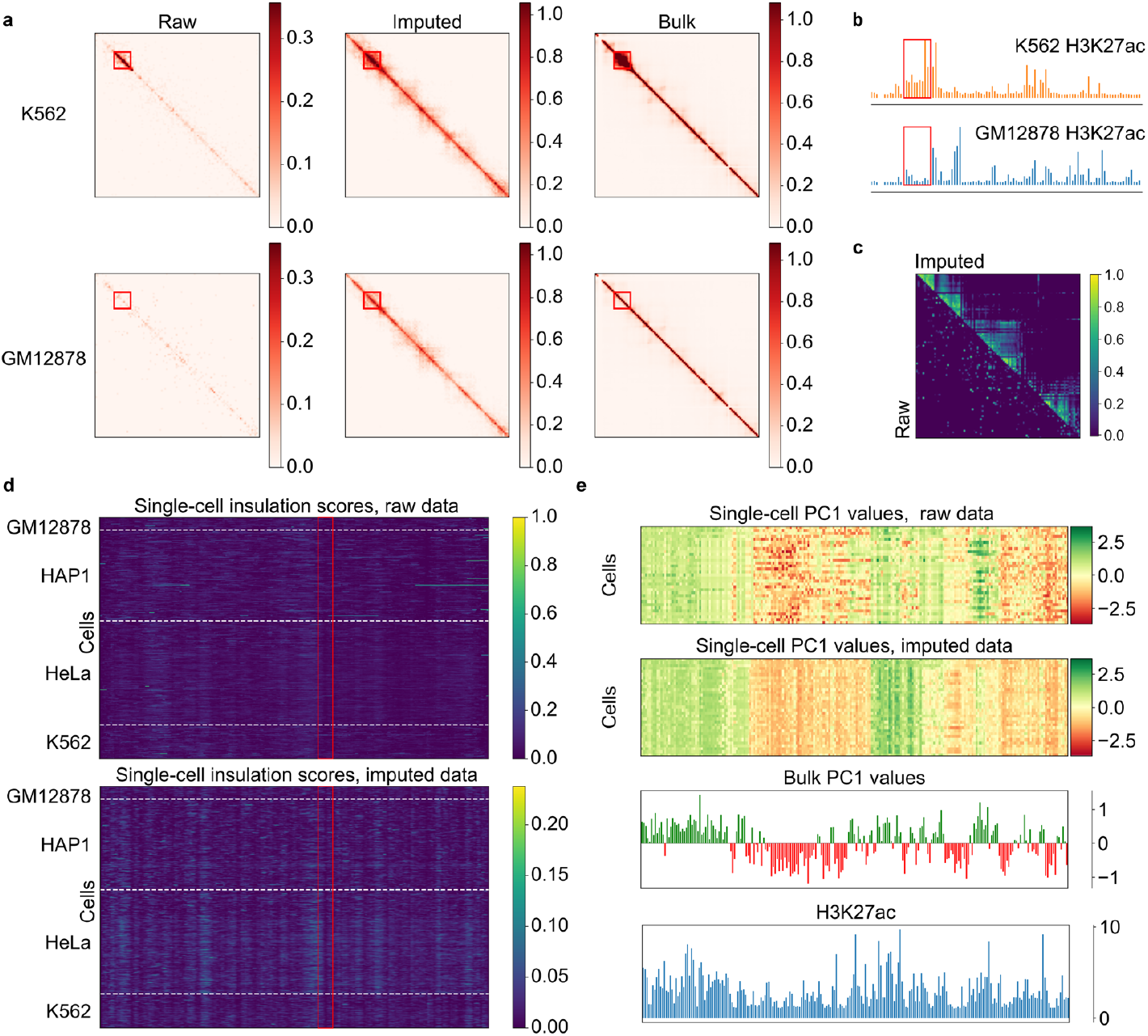
Multi-scale chromatin structure analysis from raw and imputed scHi-C data using Hi-Cformer. **a**, Averaged contact maps at 50-kb resolution for K562 and GM12878 cells (top and bottom rows) in the chr9:132,950,000–138,000,000, shown as raw single-cell data (left), imputed data using Hi-Cformer (middle), and corresponding bulk Hi-C data (right). Red boxes highlight a subregion (chr9:133,550,000–134,050,000) around the ABL1 locus. **b**, ChIP-seq tracks of H3K27ac in GM12878 and K562 across the same region as in **a**, with the red box indicating the same subregion around the ABL1 locus. **c**, Pearson correlation matrix computed from pseudo-bulk contact maps (chr9:132,950,000–138,000,000, 50-kb resolution) of GM12878 cells, obtained by averaging single-cell contact maps from Ramani2017 dataset. Each entry reflects the Pearson correlation between rows of the contact map. Lower and upper triangles show results before and after imputation, respectively. **d**, Single-cell insulation scores in chr9:100,000,000–140,000,000 at 50-kb resolution for four cell types (GM12878, HAP1, HeLa, and K562), computed from raw (top) and imputed (bottom) data. Red boxes indicate example TAD-like boundaries. **e**, Single-cell values of the first principal component (PC1) representing A/B compartment scores for GM12878 of chr1 at 1-Mb resolution. From top to bottom: PC1 values derived from raw single-cell data, PC1 values from imputed single-cell data, bulk-level PC1 values, and H3K27ac ChIP-seq signals. All tracks are shown over the same genomic region at the same resolution for direct comparison.

We further explored the impact of imputation on the identification of structural features in scHi-C data, including TAD-like domains and A/B compartments. TADs are regions of high chromatin contact frequency observed in bulk Hi-C contact maps^11^. Recent studies have reported similar structures, TAD-like domains, in scHi-C contact maps^40^. We extracted scHi-C contact maps for the chr9:132,950,000–138,000,000 at 50-kb resolution, both before and after imputation, from the Ramani2017 dataset. We then performed TAD-like domain boundary calling using the method in Higashi^27^ that identifies local minima of single-cell insulation scores as domain boundaries. The imputed data reveal clearer TAD-like domain boundaries than the raw data (Fig. 4d and Supplementary Fig. S11). By organizing single-cell insulation scores by cell type, we observed certain TAD-like domain boundaries that appear to be cell type-specific. For instance, in the region highlighted by the red box in Fig. 4d, the single-cell insulation scores of HeLa cells display clearer TAD-like boundaries relative to those from other cell types.

A/B compartments reflect the segregation of the genome into transcriptionally active and inactive regions at the mega-base scale^5^. We sought to explore how imputation influences the identification of A/B compartments. We extracted scHi-C contact maps of chr1 at 1-Mb resolution, both before and after imputation, from the two cell types (K562 and GM12878) in the Ramani2017 dataset. Then we conducted A/B compartment calling using the method in Higashi^27^, which generates comparable continuous A/B compartment scores by applying PCA to the pooled Pearson correlation matrix and projecting individual single-cell matrices using the resulting PCA components. Compared with the A/B compartment annotations derived from raw single-cell data, the compartment scores obtained from imputed data are more consistent with those from pooled pseudo-bulk data. To assess the biological relevance of the compartment scores, we examined their correspondence with H3K27ac ChIP-seq signals^41^. Regions enriched for H3K27ac tend to display scores with consistent polarity—either predominantly positive or negative—supporting the view that the scores reflect meaningful patterns of chromatin activity and compartmentalization (Fig. 4e and Supplementary Fig. S12). By considering compartment scores as cell embeddings, clustering evaluation and visualization further demonstrate that the compartment scores derived from imputed data exhibit stronger cell-type specificity (Supplementary Fig. S13).

### Supervised cell type annotation of scHi-C data with Hi-Cformer

As a versatile framework, Hi-Cformer can also be adapted for supervised cell type annotation, leveraging the existing labeled scHi-C data to directly identify cell types without relying on unsupervised clustering, thus avoiding the cumbersome process of manual annotation. This adaptation involves incorporating a simple discriminator on top of the low-dimensional cell embeddings, with the addition of a classification loss function during training. We compared Hi-Cformer with three supervised cell type annotation methods from Zhou et al.^30^: scHiClassifier, logistic regression (LR), and random forest (RF). For all baselines, we followed the same feature extraction and training procedures as described in their original study.

Evaluation was first conducted on five scHi-C datasets using five-fold cross validation. As shown in Fig. 5a, Hi-Cformer achieves consistent performance advantages across datasets, with overall average improvements of 6.24% in accuracy, 8.03% in Cohen’s kappa, and 14.00% in macro-F1 compared with the second-best method on each dataset. These metrics capture different facets of annotation quality: accuracy reflects overall correctness, kappa accounts for chance agreement across imbalanced cell populations, and macro-F1 is particularly sensitive to performance on minority or hard-to-distinguish cell types. The magnitude of improvement varies across datasets due to differences in data complexity, the number of cell types, and the separability of chromatin interaction patterns. The largest performance gain appears in the Lee2019 dataset, where Hi-Cformer achieves a markedly higher macro-F1 than all baselines. This dataset contains numerous neuronal subtypes with subtle chromatin interaction differences, making macro-F1 a stringent indicator of a model’s ability to separate fine-grained cell populations. The improvement underscores Hi-Cformer’s ability to resolve subtle chromatin differences that baseline methods miss (Fig. 5b), where it separates closely related neuronal subtypes—such as L2/3, L4, L5, and L6—that other methods largely collapse into the same class. Hi-Cformer also demonstrates superior performance in identifying rare cell types. In the Tan2021A dataset, the medium spiny neuron, a rare cell type comprising only 37 cells, is effectively recognized by Hi-Cformer. In contrast, RF fails to annotate this cell type, while scHiClassifier misannotates many other cell types as medium spiny neurons (Fig. 5c).

**Fig. 5.**
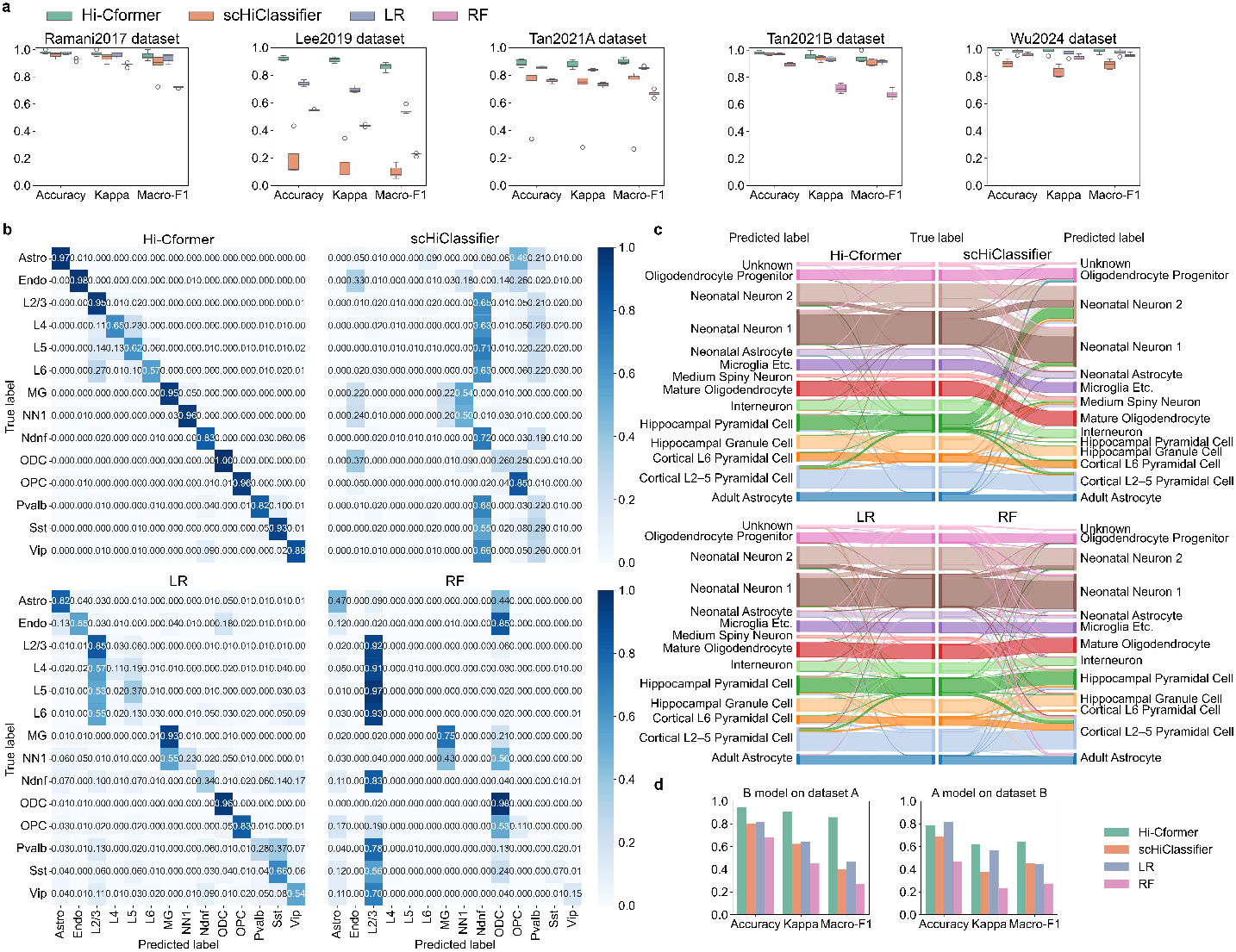
Comparison of Hi-Cformer with baseline methods for supervised cell type annotation on scHi-C data. Hi-Cformer, scHiClassifier, logistic regression (LR), and random forest (RF) were evaluated for cell type annotation performance across five public scHi-C datasets. **a**, Five-fold cross validation results on the Ramani2017, Lee2019, Tan2021A, Tan2021B, and Wu2024 datasets are shown, evaluated using accuracy, Cohen’s kappa, and macro-F1 score. The upper, middle, and lower edges of the boxes denote the upper quartile, median, and lower quartile, respectively, and error bars denote the maximum and minimum values, excluding outliers. **b**, Confusion matrix derived from different methods on the Lee2019 dataset. The results are aggregated from all validation folds in a five-fold cross validation. Each value in the matrix represents the fraction of samples predicted for a given cell type among all samples of the true cell type. **c**, Sankey plot for illustrating annotated results on the Tan2021A dataset for each cell type. Ground truth labels are shown in the center, and predicted cell types are shown on the left and right. **d**, Inter-dataset annotation performance of Hi-Cformer and baseline methods between the Tan2021A and Tan2021B datasets, evaluated using accuracy, Cohen’s kappa, and macro-F1 score.

These results highlight the effectiveness of Hi-Cformer in intra-dataset annotation tasks. To further assess these methods under more practical and challenging conditions, we evaluated their performance in inter-dataset cell type annotation, where batch effects such as technical variations may affect annotation accuracy. Tan2021A dataset consists of cells from the cortex and the hippocampus across the first postnatal year, while Tan2021B dataset contains the visual cortex of dark-reared and control mice during the first month. The biological and experimental differences between these datasets pose significant challenges for inter-dataset evaluation. After unifying the label granularity between the two datasets (details in Methods), we trained Hi-Cformer and baseline methods on one dataset and evaluated them on the other. Hi-Cformer maintains clear performance advantages in these inter-dataset settings (Fig. 5d), demonstrating adaptability across datasets. Together, these findings suggest that Hi-Cformer not only learns robust cell representations but also holds promise as a reliable framework for cell-type annotation in future scHi-C studies.

## Discussion

In this study, we present Hi-Cformer, a transformer-based framework for analyzing scHi-C data. Hi-Cformer generates embeddings that effectively capture cell-type-specific chromatin patterns, facilitating clearer distinctions among diverse cell populations. Its imputation robustly recovers missing contacts in scHi-C data, improving data quality and revealing cell type-specific regions. The variant of Hi-Cformer leverages scHi-C features for accurate cell type annotation, underscoring its potential for comprehensive single-cell chromatin conformation analysis.

The key innovation of Hi-Cformer lies in its multi-scale transformer architecture, which models chromatin contact patterns across different scales while maintaining single-cell resolution. The model captures both fine-grained and overall chromatin interaction patterns within individual cells. The chromosome-aware attention mechanism enables Hi-Cformer to capture both intra- and inter-chromosomal dependencies, integrating long-range structural relationships in a data-driven manner without relying on predefined neighborhoods or fixed-scale assumptions. Furthermore, the design of Hi-Cformer offers practical flexibility, permitting straightforward modifications to incorporate analysis of regions of interest at different resolutions and to support downstream tasks such as cell type annotation with relative ease.

There are several potential directions to further improve Hi-Cformer. First, as a transformer-based model for scHi-C data, Hi-Cformer represents input contact maps as sequences of token-like embeddings. This token-based formulation, along with additional embeddings, provides a flexible basis for incorporating other omics modalities. For example, DNA methylation or gene expression data could be introduced through newly designed embeddings, allowing joint modeling in a unified framework^16,42^. Second, potential extensions of Hi-Cformer also include the integration of inter-chromosomal contacts and the modeling of higher-order multi-way interactions to more comprehensively capture the complexity of 3D genome organization. Incorporating inter-chromosomal contacts enables the representation of spatial relationships across chromosomes, which are frequently absent in current models. The inclusion of multi-way interactions facilitates the characterization of chromatin structures that extend beyond pairwise contacts. These extensions may be implemented within the Hi-Cformer framework through the development of new encoding and decoding modules. Collectively, such advancements have the potential to improve the analysis of scHi-C data, offering new opportunities for deepening our understanding of 3D genome organization. Finally, Hi-Cformer, as a transformer-based model trained with a self-supervised objective using MLM, aligns with recent trends in foundation models^43,44^, offering a scalable framework for learning general-purpose representations of single-cell 3D genome organization. With sufficient training on large and diverse datasets, such models could support transfer learning across biological conditions and tasks, enabling various downstream applications.

## Methods

### Model architecture of Hi-Cformer

Hi-Cformer is designed as a unified multi-scale transformer-based method that learns 3D genome organization from sparse scHi-C contact maps. The model consists of three major modules: a multi-scale encoder module that extracts chromatin features spanning chromosome-level to local genomic contexts, a transformer module that integrates these features using biologically informed attention constraints, and a multi-scale decoder module that reconstructs chromatin contact patterns at multiple organizational scales. Together, these modules enable Hi-Cformer to capture hierarchical chromatin structures while maintaining the native organization of each chromosome.

#### Multi-scale encoder module

scHi-C experiments generate, for each cell, a collection of paired genomic loci that represent putative chromatin contacts. For computational analysis, these contact pairs are typically aggregated into chromosome-wise two-dimensional contact matrices, where each entry encodes the frequency of contacts between two genomic bins at a specified resolution (for example, 1-Mb bin size). For a given cell, Hi-Cformer operates on a set of intra-chromosomal contact maps defined as:

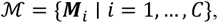

where each chromosome *i* has a contact map 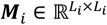 with resolution fixed across chromosomes but matrix sizes varying due to chromosome length differences. To preserve their native shapes, the maps in ℳ are processed as an ordered list rather than being reshaped into a common dimension.

The multi-scale encoder extracts two complementary levels of chromatin features: (1) chromosome-level representations that capture genome organization of entire chromosomes, referred to as chromosomal map embeddings, and (2) block-level representations that capture local chromatin structures across multiple genomic scales, referred to as block embeddings.

To obtain fixed-dimensional embeddings for entire chromosomes, we perform PCA separately for each chromosome using all cells in the training dataset. Specifically, for chromosome *i*, we first flatten each contact map from cell *j* into a vector

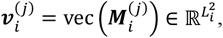

and collect these vectors across all cells to form a data matrix

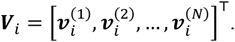

PCA is performed on ***V***_*i*_, and the top *d* principal components are used to embed each chromosome into a fixed-dimensional representation:

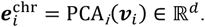

This procedure yields chromosome-level embeddings that are invariant to chromosome length and capture major axes of variation in large-scale chromatin architecture.

Local chromatin features are extracted by applying a set of multi-scale encoders to diagonal blocks of each contact map. For block size *s*_*l*_ ∈ {8,16,32,64,128}, chromosome *i* is partitioned into 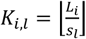 non-overlapping diagonal blocks 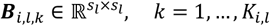. Each block is embedded using a scale-specific MLP encoder with output dimension *d*. For block ***B***_*i,l,k*_,

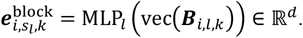

This design enables Hi-Cformer to capture local chromatin features spanning multiple genomic scales.

All embeddings—chromosomal map embeddings and block embeddings—are concatenated into a single ordered sequence, arranged first by chromosome index, then by block scale, and finally by block position: 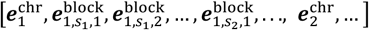 to form the embedding sequence ***E*** ∈ ℝ^*n*×*d*^, where *n* denotes the total number of embeddings. By default, the model operates on scHi-C contact maps at 1-Mb resolution, with the embedding dimension *d* = 128. This ordered embedding sequence serves as the input to the transformer module, enabling Hi-Cformer to model chromatin context across chromosomes and scales in a manner analogous to language models processing token sequences.

#### Transformer module

Because the embeddings in ***E*** come from distinct chromosomes, scales, and positions, the transformer requires contextual identifiers analogous to positional embeddings in natural language processing (NLP) models. To provide this context, we introduce chromosome, scale, and position embeddings, each capturing a different structural attribute of chromatin.

Let the embedding dimension be *d*. We designed three types of trainable information embeddings: (1) Chromosome embeddings {***c***_*i*_ ∈ ℝ^*d*^ ∣ *i* = 1, 2, …, *C*}, marking the chromosome from which each embedding originates. (2) Size embeddings {***s***_*i*_ ∈ ℝ^*d*^ ∣ *i* = 1,2, …, *S*}, distinguishing blocks of different genomic scales. (3) Position embeddings {***P***_*i*_ ∈ ℝ^*d*^ ∣ *i* = 1,2, …, *K*_*max*_}, where *K*_*max*_ denotes the maximum number of blocks produced by any chromosome at the smallest block size. We use this upper bound to ensure that positional indices can accommodate the longest possible block sequence across chromosomes. The information embeddings are added to the original embedding sequence, resulting in a new embedding sequence 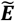, whose elements are computed as follows:

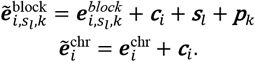

To integrate these embeddings, Hi-Cformer employs a chromosome-aware transformer encoder with a biologically constrained attention mechanism, specifically tailored to 3D genome organization. By default, the transformer module consists of four stacked transformer encoder blocks. Each block operates on inputs and outputs of dimension *d* and uses 8 attention heads. Let 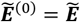 denote the input to the transformer. The output of the *l*-th encoder block is denoted by 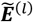 and is computed as

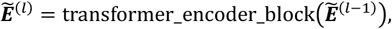

where transformer_encoder_block(·) is a single transformer block and *l* = 1, …, *L* with *L*=4 in our default configuration.

A central innovation of Hi-Cformer is its biologically informed self-attention, which explicitly encodes constraints on which chromatin regions are allowed to interact. Conceptually, the attention mechanism serves two purposes: (1) to model local and long-range 3D contacts within each chromosome via block-level interactions, and (2) to propagate global context between chromosomes via chromosome-level embeddings. To achieve this, we redesign the self-attention with a structured attention mask that respects the hierarchical organization of the genome. For the *l*-th transformer block, we first compute the Query, Key and Value matrices from 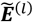:

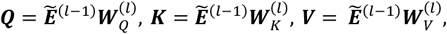

Where 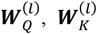, and 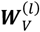 are trainable weight matrices used to compute the Query (***Q***), Key (***K***), and Value (***V***), respectively, and *d*_*k*_ denotes the dimension of the key vector. The biologically constrained attention is then defined as

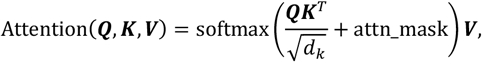

where attn_mask ∈ ℝ^*n*×*n*^ encodes which pairs of embeddings are allowed to attend to each other.

To reflect biological constraints on chromatin organization, we design attn_mask with the following structured pattern. Block embeddings are permitted to attend only to embeddings from the same chromosome—namely, other block embeddings and the corresponding chromosomal map embedding. In contrast, chromosomal map embeddings are allowed to attend globally to all chromosome-level embeddings across the genome, while still attending to block embeddings within their own chromosome. Formally, for the *i*-th query embedding and the *j*-th key embedding, we define

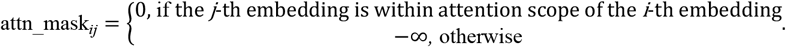

By adding this mask before the softmax operation, we zero out disallowed cross-chromosome block-block interactions, while preserving global chromosome-level communication through the chromosome embeddings. This structured attention mask is a key component that distinguishes Hi-Cformer from standard transformers and embeds biological priors directly into the attention pattern.

The final output of the transformer module is denoted by 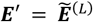. The contextual chromosomal map embeddings in ***E***′ after the transformer module can be seen as containing not only all the information of their respective chromosomes but also information from other chromosomes. And each block embedding not only contains information about its corresponding contact region but also includes additional information from other regions within the same chromosome.

#### Multi-scale decoder module

The multi-scale decoder module is designed to reconstruct chromatin contact patterns at the cell, chromosome, and block levels, thereby enabling Hi-Cformer to learn a unified latent representation of 3D genome organization across scales. After the transformer generates contextual embeddings, the decoder processes these embeddings through three parallel pathways, each dedicated to a specific structural resolution.

To obtain a compact representation of the overall chromatin architecture of a cell, we concatenate all contextual chromosomal map embeddings into a matrix ***E***′^map^ ∈ ℝ^*C*×*d*^ from ***E***′. A two-layer fully connected network maps this matrix to a low-dimensional cell embedding. Specifically, the first layer reduces flattened ***E***′^map^ to a 64-dimensional vector and outputs the cell embedding ***e***′^cell^ ∈ ℝ^*g*^, where *g* = 64. The second layer takes ***e***′^cell^ as input and outputs ***r***^cell^ ∈ ℝ^*o*^. The dimensionality *o* is defined to exactly match the length obtained by concatenating and flattening the upper-triangular regions of all intra-chromosomal contact maps. Because scHi-C contact matrices are symmetric, the upper-triangular portion uniquely specifies the full matrix; therefore, aligning ***r***^cell^ to this dimensionality enables the decoder to directly generate a complete, cell-level reconstruction of chromatin interactions. Consequently, ***r***^cell^ serves as the imputation result for the cell.

To reconstruct chromosome-wise contact maps, each contextual chromosomal map embedding 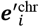 is passed through a single fully connected layer that outputs a square matrix 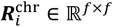, where *f* is set to 112 in this work. The reconstructed signals from the chromosome-level decoder are considered scaled versions of the chromosomal contact maps. Therefore, the original frequency maps are interpolated to the specified size before computing the reconstruction loss.

The block-level decoder consists of several single-layer fully connected neural networks, taking an input 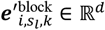 and producing an output 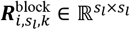 to reconstruct the corresponding block.

The purpose of these decoders is to train the model through reconstruction tasks, thereby obtaining low-dimensional representations of multi-scale structures, including cells, chromosomes, and blocks.

### Model training

To optimize multiscale representation learning, the model is trained using reconstruction objectives defined at three hierarchical levels: cell, chromosome, and block. Each loss term serves a distinct purpose aligned with the corresponding biological and structural scale. The cell-level loss integrates information across all chromosomes and drives the learning of a global cell embedding. The chromosome-level loss supervises the reconstruction of individual chromosomal contact maps, thereby shaping the chromosomal map embeddings. The block-level loss focuses on recovering fine-scale structural units and thus specifically trains the block embeddings. The final training objective is defined as the weighted combination of these three reconstruction losses. Assuming that 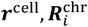, and 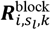 correspond to the original signals 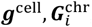, and 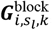, respectively, the training loss function is computed as follows:

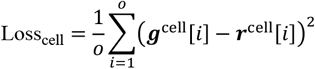

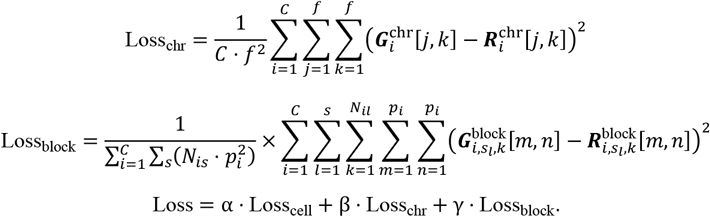

Here, α, β, and γ are tunable hyperparameters. Prior to the main training, Hi-Cformer undergoes a phase referred to as preheating. During preheating, the transformer module is removed, while other modules and loss functions remain unchanged. The only modification is that the decoder receives 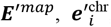, and 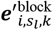 extracted from ***E***′ instead of 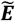. The decoder then produces 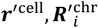, and 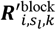, which replace 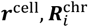, and 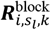 in the loss computation, allowing the calculation of the same losses as described above.

The application of MLM tasks in the BERT model^32^ has shown remarkable efficiency and brought profound changes to the field of NLP. Inspired by these masked language modeling tasks, this work also designs a masked language modeling task to train the model. Specifically, given a masking ratio, after obtaining the embedding sequence ***E***, all embeddings (including both chromosomal map and block embeddings) belonging to the same chromosome are randomly replaced with a learnable special embedding [MASK] at the specified ratio. The modified embedding sequence ***E*** is then used as the input to the transformer module. In our experiments, the default masking ratio is set to 20%. For highly sparse contact maps, we recommend increasing the masking ratio to encourage more effective learning of missing or noisy regions. For less sparse data, a lower masking ratio may be preferable to preserve more of the original structural information during training.

### Data processing

In this work, we utilized several publicly available scHi-C datasets with cell type labels, as well as bulk Hi-C and ChIP-seq datasets for reference analyses. The scHi-C datasets include the Ramani2017 dataset^15^ consisting of four human cell lines (GM12878, HAP1, HeLa, and K562); the Lee2019 dataset^16^ from human prefrontal cortex cells; the Tan2021A and Tan2021B datasets^17^ from mouse brain development; and the Wu2024 dataset^18^ comprising four human cell lines (BJ, GM12878, K562, and eHAP). For bulk Hi-C reference, we used contact maps for the K562 and GM12878 cell lines from the Rao2014 dataset^14^. In addition, we obtained ChIP-seq tracks of H3K27ac in GM12878 and K562 generated from the ENCODE project^41^ to support downstream validation and biological interpretation.

Following the data processing pipeline of Higashi^27^, cell filtering was performed on all datasets, excluding cells with fewer than 2,000 total reads or a genome span smaller than 500 kb. After preprocessing, the Ramani2017 dataset consists of 620 cells from four human cell types, the Lee2019 dataset consists of 4,238 cells from 14 human cell types, the Tan2021A dataset consists of 1,692 cells from seven mouse cell types, the Tan2021B dataset consists of 1,954 cells from 14 mouse cell types, and the Wu2024 dataset consists of 431 cells from four human cell types.

The processed read pairs were binned into contact matrices at a specific genomic resolution. In this work, we only retained contact pairs within the same chromosome. The raw read data for all chromosomes of a cell was transformed into symmetric matrices of varying sizes, which served as the input for the model. For datasets with significant batch effects, such as the Lee2019 dataset, we applied BandNorm suggested by Zheng et al.^28^ to the data. In the dropout experiment, we randomly discarded raw read pairs with a certain probability, and then performed the processing steps mentioned above.

### Downstream analysis

#### Unsupervised feature extraction

We evaluated the low-dimensional cell embeddings generated by Hi-Cformer on five publicly available scHi-C datasets with annotated cell types, all processed at 1-Mb resolution. To benchmark the performance, we compared Hi-Cformer against six established baseline methods: Higashi^27^, scDEC-Hi-C^29^, HiCRep/MDS^26^, scHiCluster^23^, PCA^34^, and LDA^24^. Each baseline method was implemented following its respective publication and official codebase, using either recommended or default hyperparameter settings to ensure fairness. For Higashi, we used the version corresponding to the commits on October 11, 2021, from its GitHub repository. To ensure comparability, the dimensionality of the embeddings used for evaluation was set to 64 across all methods, except for scDEC-Hi-C. Due to the instability of its GAN-based architecture, we were unable to obtain a converged model in its second-stage training under default settings. Therefore, for scDEC-Hi-C, we used the concatenated chromosome-wise embeddings generated from its first-stage autoencoder as the final cell representation, resulting in a higher-dimensional embedding of 1150 dimensions.

For each method, we applied Leiden clustering to assess their ability to capture cell type heterogeneity. To align the number of clusters with the ground-truth cell type annotations, we employed a binary search strategy to select the clustering resolution. The performance was quantified using NMI, ARI and cLISI. The detailed formulas for computing these metrics are available in Supplementary Note A.1.

To assess the robustness of Hi-Cformer to data sparsity, we simulated dropout in the Ramani2017 dataset by applying random contact removal directly to the original paired-end read files. Each contact was independently retained with a fixed probability 1 − *p*, where *p* is the dropout rate. The resulting contact pairs were then aggregated into single-cell contact frequency matrices at 1-Mb resolution, following the same preprocessing pipeline as in the main analysis. All model settings, training procedures and evaluation criteria remained consistent with those used in the original embedding experiment on the Ramani2017 dataset, ensuring that the robustness evaluation focused solely on the effect of data sparsity.

To evaluate the contribution of each component of Hi-Cformer, we conducted ablation experiments by removing the transformer module, preheating strategy, and MLM task. Three variants of the model were created by sequentially omitting these components, and the embeddings generated by each model were compared to those of the original Hi-Cformer model using the Ramani2017 dataset.

#### Data imputation

To evaluate the ability of Hi-Cformer to impute scHi-C contact maps, we compared it with two other imputation methods, Higashi^27^ and scHiCluster^23^. Unless otherwise specified, the model parameters for Hi-Cformer were identical to those used in the cell embedding evaluations. For each cell, the reconstructed signal output from Hi-Cformer, denoted as ***r***^cell^, was regarded as the imputed contact map.

To quantify similarity between the imputed contact maps and reference contact maps, we computed both the Pearson correlation coefficient (PCC) and cosine similarity. These metrics were calculated between (1) the imputed single-cell contact maps and the corresponding bulk Hi-C maps, and (2) the imputed single-cell maps and pseudo-bulk maps obtained by averaging scHi-C contact maps from cells of the same type. To avoid redundancy, only the upper triangular part of each contact map was extracted and flattened into a one-dimensional vector before similarity computation.

For the dropout experiments, we generated corrupted datasets with varying dropout ratios using the same procedure as in the cell embedding evaluation, treating the raw scHi-C data as the ground truth. Imputation performance was assessed using PCC, cosine similarity, the structural similarity index measure (SSIM) and the peak signal-to-noise ratio (PSNR). The exact formulas for these four metrics are provided in Supplementary Note A.2. To further evaluate how well the imputed contact maps preserve cell-type heterogeneity, we applied PCA to reduce each imputed map to a 64-dimensional representation, followed by clustering on these low-dimensional embeddings.

In the section “Hi-Cformer facilitates the identification of cell-type-specific structures”, we introduced a specialized 50-kb resolution region (chr9: 100,000,000–140,000,000) as input. Specifically, this region was fed into an additional fully connected layer to transform it into a 128-dimensional embedding. To distinguish this region from others, a unique size embedding was added to it. In the attention mechanism, the embedding of this special region was configured to calculate the attention score with its corresponding chromosomal map embedding, while preventing other embeddings from attending to it. Finally, this specialized embedding passed through the transformer module and was processed by an added deconvolutional network to reconstruct the signal. The reconstructed signal, treated as the imputation result, was compared to the original input to compute the Mean Squared Error (MSE) loss, which was incorporated into the Loss_*block*_ for joint optimization.

For visualization of correlation patterns, we first applied the observed-expected normalization procedure described by Lieberman-Aiden et al.^5^ to the contact maps and then computed the Pearson correlation matrix. The methods for calling TAD-like boundaries and A/B compartments followed the approach proposed in Higashi.

#### Supervised cell type annotation

To enable supervised cell type annotation, we extended Hi-Cformer by attaching a single fully connected layer to the cell embedding. The output of this layer was trained with a cross-entropy loss, which was added to the original Hi-Cformer objective during optimization.

We compared Hi-Cformer with three baseline methods: scHiClassifier^30^, logistic regression (LR), and random forest (RF). For LR and RF, we used the four feature sets extracted from scHi-C contact maps as proposed by Zhou et al.^30^ All methods were evaluated on five datasets using five-fold cross validation. Cell type annotation performance was assessed using three metrics: accuracy, Cohen’s Kappa, and macro-F1, with details of metric computation provided in Supplementary Note A.3.

For cross-dataset evaluation between the Tan2021A and Tan2021B datasets, we performed label alignment to enable meaningful comparison. Specifically, in the evaluation phase, several neuronal subtypes in Tan2021A were merged into a single “Neuron” category (see Supplementary Note A.4 for the exact cell types), while the original labels were retained during training. This means that when a model trained on Tan2021A dataset is applied to Tan2021B dataset, the neuron subtype prediction (such as Neonatal Neuron 1) it gives will be treated as the “Neuron” class. When a model trained on the Tan2021B dataset is applied to the Tan2021A dataset, its prediction is considered correct if it annotates a neuronal subtype as the general “Neuron” category. To ensure fair comparison, cell types that were not present in the training set were excluded from metric calculation in the test set.

## Data availability

We used the following publicly available datasets: scHi-C data from Ramani et al.^15^ (GEO: GSE84920); sn-m3c-seq from Lee et al.^16^ (GEO: GSE130711); scHi-C data (Dip-C) from Tan et al.^17^ (GEO: GSE164203 and GEO: GSE146397); scHi-C data from Wu et al.^18^ (GEO: GSE240128); bulk Hi-C data of K562 and GM12878 from Rao et al.^14^ (GEO: GSE63525); and ChIP-seq of H3K27ac in GM12878 and K562 from ENCODE^41^ (GEO: GSE29611).

## Code availability

Hi-Cformer is freely available on GitHub (https://github.com/Xiaoqing-Wu02/Hi-Cformer).

## Acknowledgements

This work was supported by the National Key Research and Development Program of China (grant nos. 2025YFC3409300, 2023YFF1204802), the National Natural Science Foundation of China (grant nos. 32550616, 62273194) and Beijing Natural Science Foundation (grant no. L242026).

## Author contributions

R.J. conceived the study and supervised the project. X.W. and X.C. designed, implemented and validated Hi-Cformer. X.W., X.C., and R.J. wrote the manuscript, with input from all the authors.

## Competing interests

The authors declare no competing interests.

## Notes

### Competing Interest Statement

The authors have declared no competing interest.

### Summary of Updates

Figure 1 has been revised to improve clarity. No other major changes were made.

